# Controlled bio-inspired self-organised criticality

**DOI:** 10.1101/2021.05.05.442730

**Authors:** Tjeerd V. olde Scheper

## Abstract

The control of extensive complex biological systems is considered to depend on feedback mechanisms. Reduced systems modelling has been effective to describe these mechanisms, but this approach does not sufficiently encompass the required complexity that is needed to understand how localised control in a biological system can provide global stable states. Self-Organised Criticality (SOC) is a characteristic property of locally interacting physical systems which readily emerges from changes to its dynamic state due to small nonlinear perturbations. Small changes in the local states, or in local interactions, can greatly affect the total system state of critical systems. It has long been conjectured that SOC is cardinal to biological systems that show similar critical dynamics and also may exhibit near power-law relations. Rate Control of Chaos (RCC) provides a suitable robust mechanism to generate SOC systems which operates at the edge of chaos. The bio-inspired RCC method requires only local instantaneous knowledge of some of the variables of the system, and is capable of adapting to local perturbations. Importantly, connected RCC controlled oscillators can maintain global multi-stable states, and domains with power-law relations may emerge. The network of oscillators deterministically stabilises into different orbits for different perturbations and the relation between the perturbation and amplitude can show exponential and power-law correlations. This is representative of a basic mechanism of protein production and control, that underlies complex processes such as homeostasis. Providing feedback from the global state, the total system dynamic behaviour can be boosted or reduced. Controlled SOC can provide much greater understanding of biological control mechanisms, that are based on distributed local producers, remote consumers of biological resources, with globally defined control.

**Author summary:** Using a nonlinear control method inspired by enzymatic control, which is capable of stabilising chaotic systems into periodic orbits or steady-states, it is shown that a controlled system can be created that is scale-free and in a critical state. This means that the system can easily move from one stable orbit to another using only a small local perturbation. Such a system is known as self-organised criticality, and is shown in this system to be deterministic. Using a known perturbation, it will result in a scale-free response of the system that can be in a power law relation. It has been conjectured that biosystems are in a self-organised critical state, and these models show that this is a suitable approach to allow local systems to control a global state, such as homeostatic control. The underlying principle is based on rate control of chaos, and can be used to understand how biosystems can use localised control to ensure stability at different dynamic scales without supervising mechanisms.

## Introduction

The physical universe, in which biological systems exist, is inherently nonlinear, chaotic and noisy. The concepts of dynamic stability in complex biosystems is accepted [1], but not very well understood at all levels. The capacity of biosystems to control their internal and external response to the chaotic environment can be contributed to their ability to adapt in a nonlinear manner to the highly variable environment. The facility to quickly change state, apparently scale-free, is essential to the survival of each organism [1]. Furthermore, the high level stability of a larger organism must be constructed from the low-level dynamic behaviour of many contributing elements, organs, and cellular behaviours. In particular, the local information available to each organ, cell, sub-cellular body, or other organic organisation, is severely limited with regards to the global state, and is also limited in its ability to respond in a timely and measured manner. Although there exists some robust understanding of the mechanisms involved in global stability of an organism [2], this is certainly not the case for the required complexity of local small elements collaborating in a nonlinear biological system to generate a globally stable dynamic state [3].

Self-Organised Criticality (SOC) is a distinct property of physical systems based on the local interactions of many small components each of which contributes to the global critical system [4, 5]. Depending on the field of study, there seems to exist different interpretations of the meaning of self-organised criticality [6]. The concept of *critical behaviour*, or just *criticality*, and its meaning have been neatly summarised by Watkins et al. [7]. Irrespective of alternative means of generating self-organised critical systems, the proposed mechanism within this paper is not based on a physical phenomenon as such, and focuses on the mathematical interpretation of such systems within dynamic systems theory. This is described by the ability of complex systems to reside at an *equilibrium point* or *critical point* within their parameter space, such that they will change from one state to another with only a small perturbation to the system [8]. Therefore, weak perturbations of the system from external sources may cause a state of change due to the critical point, where the system will evolve into the new stable or unstable state. The proposed mechanism of control ensures that the system remains stable, in the Lyapunov sense [9], but with uniquely different stable states.

Small interactions within the components of a SOC system are not significantly large with respect to the scale of each element, but they contribute to the global critical state due to nonlinear behaviour. These nonlinear local perturbations are usually observed as rapid transitions from one state to another of the global system. A typical illustration is an avalanche of snow or sand, where a previously apparent stable state rapidly changes to a new stable state. Similar critical dynamics have been determined in biological systems, at low [10] and high levels [11], and in particular in neural dynamics [12,13]. The functional role of criticality, the self-organised and self-sustaining multi-stable state of the system, appears to be mostly related to network complexity, and power-law relations within those networks [14]. However, it is still unclear if power-law relations are a true property of large scale complex systems over the entire domain of scale [15]. Furthermore, as will be shown, the emergent power-law relations may be considered only an epiphenomenon of the combined response of the nonlinear oscillators, and may exist for only part of the parameter domain, where the essential property of the system is stable periodic behaviour throughout these perturbations. For the described systems below, the means by which the criticality may emerge is not based on the separation of timescales, but on the local adjustments of the control mechanism in combination with nonlinear connectivity. The method contains all three key features for a system to be in a critical state, namely non-trivial scaling due to the external perturbation, spatiotemporal power-law correlation in respect to the total behaviour of the network, and self-tuning to the critical point where the network self-selects the periodic orbit [7].

Considerable effort has been put into deciding the characteristic properties of large and complex systems. Aspects, such as connectivity, and dynamics are known to be important, on which the principle of Artificial Neural Networks is based. Here, it is argued that the problem can be said to lie in determining how a local system, usually at a much smaller scale, can contribute to a global, stable system of which they form only a small part. Within dynamic systems theory, these local systems can act as perturbations of the overall global system. Even if the individual components are stable, the entire system may become unstable due to these perturbing elements. This is commonly seen in spatiotemporal chaos, where individual elements are dynamically stable, yet destabilise the entire system when weakly coupled [16]. Within biosystems this property, where clusters of smaller elements contribute to the total behaviour, is very common. For example, it may be seen clearly in the organs of large multi-cellular organisms [17,18]. Loss of stability in any of these localised systems may be catastrophic for the entire global system. The ability to ensure that the entire system remains stable may not depend on a specific supervisory element, but must be provided by the cooperative nature of the constituting elements. This, in effect, precludes the use of supervisory systems, even if that is a much higher order controller, such as the brain [19]. This characteristic property of biological systems to exhibit emergent dynamic behaviour is based on local small scale distributed dynamics behaviour which collaborates to produce a higher order dynamics. In particular, the ability to respond in a characteristic nonlinear manner, such as a log scale response, to input [20], as well as the regulation of producing and consuming biomatter. Different parts of the organism are involved in maintaining and developing the global system and these small conglomerates of cells or organisms can produce types of activity similar to the behaviour of larger organisations [1]. For these disparate systems to cooperate effectively, it would seem that local control is not sufficient. However, if these systems share properties with critical systems [21], it becomes feasible that by changing the response of the local system to the external conditions, a suitable state can be found that is appropriate for control to be effective at the global scale. This means that the proposed mechanism of control permits local control of homeostatic systems without external or supervisory approaches.

Homeostatic control is based on negative, as well as occasionally positive, feedback of some of the components in relation to some set point. This well known concept is enticingly simple, and has echoes of the older concepts of physical harmony, and well balanced systems. It exists in many biological systems at almost all levels, from sub-cellular molecular dynamics [22,23], through physiological phenomena (e.g. glucose and blood pressure levels) [24], neural dynamics [25], and all the way to sociological behaviour. Both from experimental data and from physiological experiences [26], this concept seems to be somewhat flawed. In dynamic systems theory it has been shown that feedback systems can only under the strictest circumstances be guaranteed to be stable. For example, it is shown that a reduced system can have multiple stable states using isolated feedback loops [27] but only for a specific parameter set. Positive and negative feedback loops may have both stabilising and destabilising effects, depending on the shape of the Jacobian matrix for only some determined stable states [28]. Lastly, even an unified control approach, emphasising the possibility of hierarchical control, avoids the issue of system stability and does not consider the effect of the complexity of each of the boxes connected within various feedback loops [2].

It has become clear that even well known homeostatic systems are very rarely stable in the dynamic sense, appearing to have multiple stable states, and do not show the properties of stable systems when perturbed. This has now been encapsulated within the concept of allostasis that permits multiple stable states within an homeostatic system [29]. Assuming that a homeostatic system is the apparent result of underlying stabilising mechanisms, rather than the mechanism itself, it may be possible to explain this concept, and its limitations in a consistent dynamic systems manner, using criticality [10,21]. Employing criticality to understand dynamic interactions allows the system to be highly variable, as indicated by multiple stable but controlled states, and at the same time be dynamically stable in the sense that each individual state does not destabilise the system. Providing dynamic stability with high variability of the emerging system will allow for a better understanding of the underlying mechanisms, and may provide possible ways to resolve the limitations of homeostatic control that currently cannot be addressed appropriately [30].

The simulations described in this paper aim to develop possible mechanisms for local control that are based on minimal models but with the capability to control indirectly the global state of complex systems. This has become feasible by means of the Rate Control of Chaos method that ensures that the dynamic system under control remains dynamically stable but allows many multiple states to co-exist within the same parameter space.

## Methods

Recently, it has been shown that nonlinear chaotic dynamics can be stabilised using the Rate Control of Chaos method. This method adjusts the rate of evolution of part of a nonlinear system such that the exponential growth of an unstable chaotic oscillator is controlled into stable limit cycles. The control is estimated using the rate of growth of some of the variables in proportion to the overall embedded phase space of those variables. This is then applied to an exponential control function that, in effect, causes the rate of change of the variable to be controlled to speed up, or slow down. The proportional rate of change is unity when no control is applied or is not changing exponentially [31,32].

This method may be regarded as an extension of the traditional biochemical enzyme control concept by adjusting the reaction rate. The ability to control the stability of biochemical reactions, by controlling the rate of reaction based only on local information, allows the biosystem to function under a wide range of conditions. Furthermore, it has been shown that this mechanism can be extended further to control higher level spatiotemporal chaos, even if the underlying dynamics of each element is chaotic in its own right [32]. The individual elements, as well as the total observable system, are stable in the sense of Lyapunov stability. This means that the system will reliably return to the same area of phase space relative to its controlled dynamics, although it does not necessarily follow that this is the same as the uncontrolled chaotic domain. Therefore, the RCC method does not eliminate entirely the chaotic properties of the underlying nonlinear system, but applies limited localised control to the system to maintain an apparently stable system. The controlled system still has many properties of the nonlinear system; it can respond nonlinearly to perturbations and can be weakly chaotic. This is illustrated in Fig 1A, where the RCC controlled system of a bienzymatic model, described below, is controlled into a stable orbit. In Fig 1B is shown the local Lyapunov estimates for the controlled system, demonstrating weak chaos. The phase space of both the controlled and uncontrolled system can be seen in Fig 1C.

**Fig 1.**
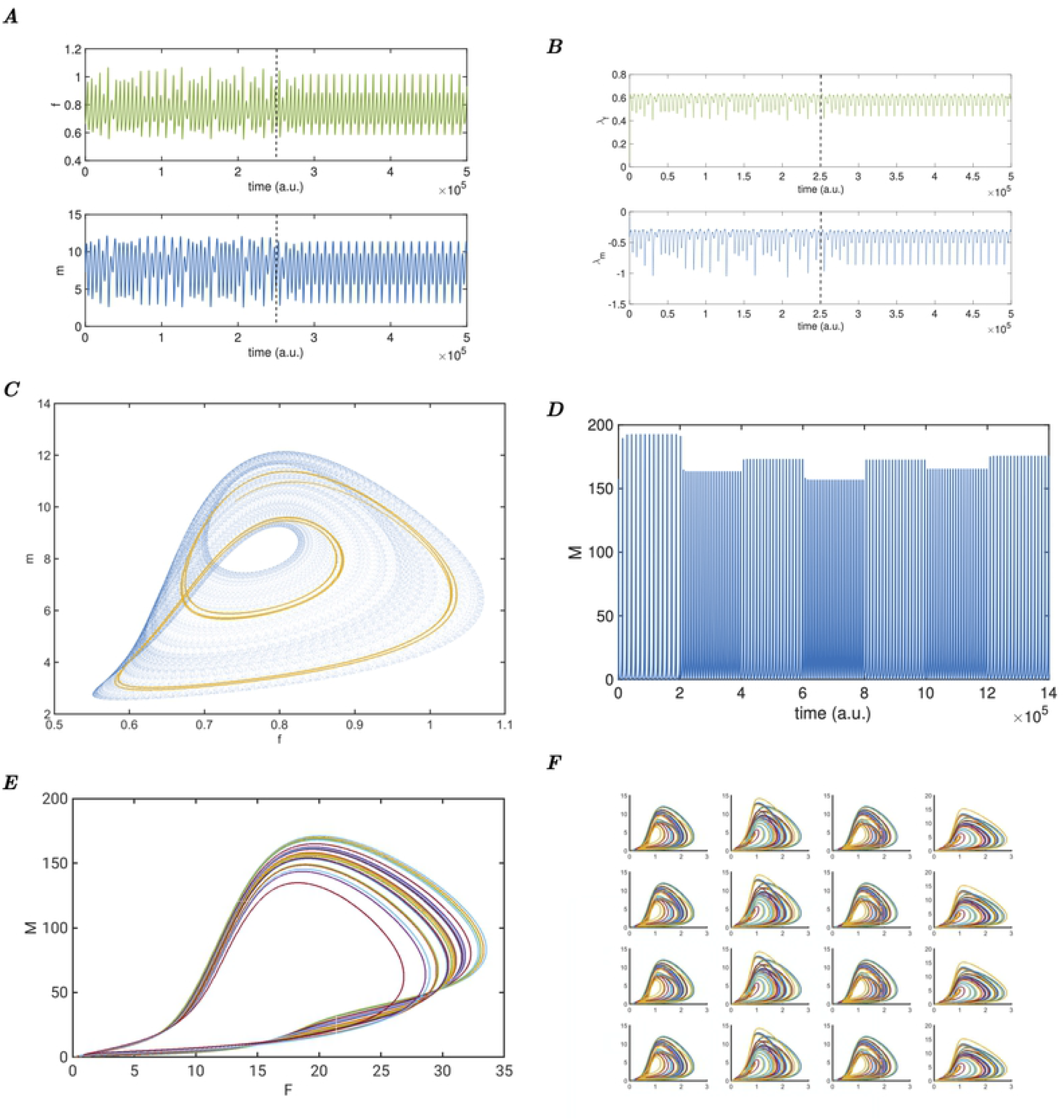
Dynamic controlled behaviour of the Berry model. ***A*** Example of the chaotic Berry model with control enabled at the dotted line, the system quickly stabilises into a two orbit. Top panel shows the modelled soluble filament *f*, the bottom panel the fixed matrix *m* in time. ***B*** Lyapunov estimates of the model in Fig 1A, showing small positive and small negative values. The control, enabled at the dotted line, does not eliminate the chaotic state, but changes the system into a stable oscillation. ***C*** Phase space plot of the chaotic model in blue, with the RCC controlled two orbit superimposed in gold, *F* versus *M*. ***D*** Total unweighted sum *M* from sixteen weakly coupled RCC controlled oscillators, with random external perturbations at fixed intervals showing 7 stable oscillations after very short transients. ***E*** Phase space plot *F* versus *M*, of 24 stable oscillations based on the total unweighted sum of the sixteen weakly coupled oscillators. Each coloured line is a single stable oscillations, as can be seen in time in Fig 1D. ***F*** Phase space plots of the sixteen systems *F* versus *M*, where each coloured line is a single stable oscillation, different due to the random external perturbations (transients removed).

The nonlinear model of a bienzymatic cycle used in this paper by Berry [33], described by (4) to (7), has been shown to be controllable using the Rate Control of Chaos (RCC) method, such that the control allows the stabilisation of the external environment by adjusting the amount of enzyme based on the local amount of one of the components *f* [34]. This model describes the two enzymes that control the formation of extracellular matrix *m* from soluble filaments f. The proteinase *p* transforms the matrix into filaments, and the transglutaminase *g* converts the filaments into matrix. Extracellular matrix is produced by neighbouring cells *r_im_* at a constant rate, and each protein decays in catalytic processes proportional to *p. r_im_* is a bifurcation parameter, that may cause the system to become unstable and chaotic. Application of Rate Control of Chaos, described by the quotient *q_f_* (1), and the two control functions *σ_p_* (2), and *σ_g_* (3), can control the production of the two enzymes *p* and *g* (in (6) and (7)) such that the system remains in controlled stable orbits for large ranges for values of the bifurcation parameter *r_im_.* Within the subsequent simulations, the model is shown as time series of the main variables *m* and *f* or phase space plots of *f* (x-axis) versus *m* (y-axis).

Furthermore, the generic properties of a two enzymatic control of a producer and consumer system, such as the formation of (extra-)cellular matrix, can be considered to be representative of many biological processes where a resource is controlled by antagonistic control using two enzymes. The model is not considered to be solely representative of this concept, but is recognisably typical for many types of control needed for biological control processes.

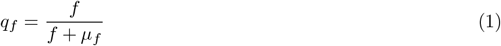

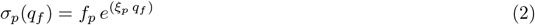

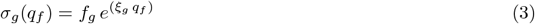

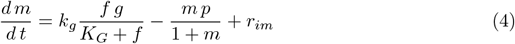

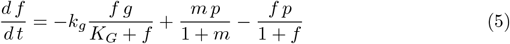

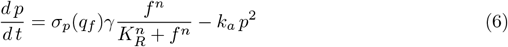

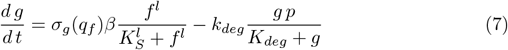

The Berry model parameters are as follows; *γ* = 0.026, *β* = 0.00075, *K_R_* = 4.5, *K_S_* = 1, *K_G_* = 0.1, *K_deg_* = 1.1, *k_g_* = *k_deg_* = 0.05, 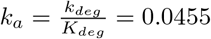 and the Hill-numbers *l* = *n* = 4. For different values of the bifurcation parameter *r_im_* in (4), the model exhibits a wide range of dynamic behaviour, including periodic cycles, bistability and chaos [33]. This parameter is kept for all oscillators within the chaotic domain. External input is provided to this parameter in the perturbation experiments (8), and this parameter is used to connect the different oscillators together using a scaled relative contribution from all the other oscillators (i.e. no self-connections):

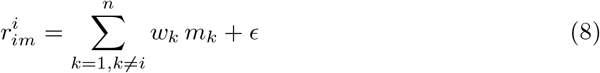

where *w_k_* the connectivity strength for the oscillator which can be either 0.00011, 0.00012, or 0.00025, and remains within the chaotic domain. *ϵ* is a uniform distributed perturbation term drawn from the domain [-1,1] and scaled to the connectivity strength of each unit. The perturbation used for the oscillators is the same random uniform distributed value applied to each oscillator, but is scaled by the randomly chosen values [7.5, 1, 8, 3.25], although each column in Fig 1F has the same perturbation value to ensure that the network is not symmetric. Furthermore, the perturbation is redrawn after a certain number of evolution steps, allowing the system to be explored for different values of perturbation, resulting in different oscillations. The connectivity strength values for each oscillator may vary, and this affects the dynamics, but not the stability. This is shown in the simulations with stepwise increase of the connectivity strength, starting from *w_k_* = [0.0001, 0.0002, 0.0003, 0.0004] for each column, and increased after 10^5^ time steps by 0.00005 (Fig 2C). The connectivities were therefore kept within the chaotic domain of the underlying oscillators, and further work is under way to demonstrate the utility of the connectivity strength within these models. The RCC method can be shown to stabilise spatiotemporal patterns, which may become unstable due to local nonlinear interaction, and is effective even when the underlying systems are chaotic [32]. The advantage of this method is that it allows nonlinear systems to be stabilised into periodic stable dynamics based only on the local dynamics of each individual oscillator. The Rate Control of Chaos parameters were also kept constant in these models, but can be varied to change the shape of the local oscillator. For the first perturbation experiments (described in figures 1-4), the control parameters in (1) are *μ_f_* = 2, and in the RCC functions for each of the enzymes *p* (2) and *g* (3) are *ξ_p_* = – 1, *f_p_* = 1, *ξ_g_* = – 1, *f_g_* = 1. In the criticality experiments with global feedback, the control is enhanced by *ξ_p_* = –3, and *ξ_g_* = –3, with no other changes. In the experiments with 64 oscillators (see figure 4), the connectivity strength for each oscillator was a randomly assigned unique value between 1 and 10. This was to show that the parameters do not need careful tuning beyond ensuring that the system is in the chaotic domain, and RCC controlled.

**Fig 2.**
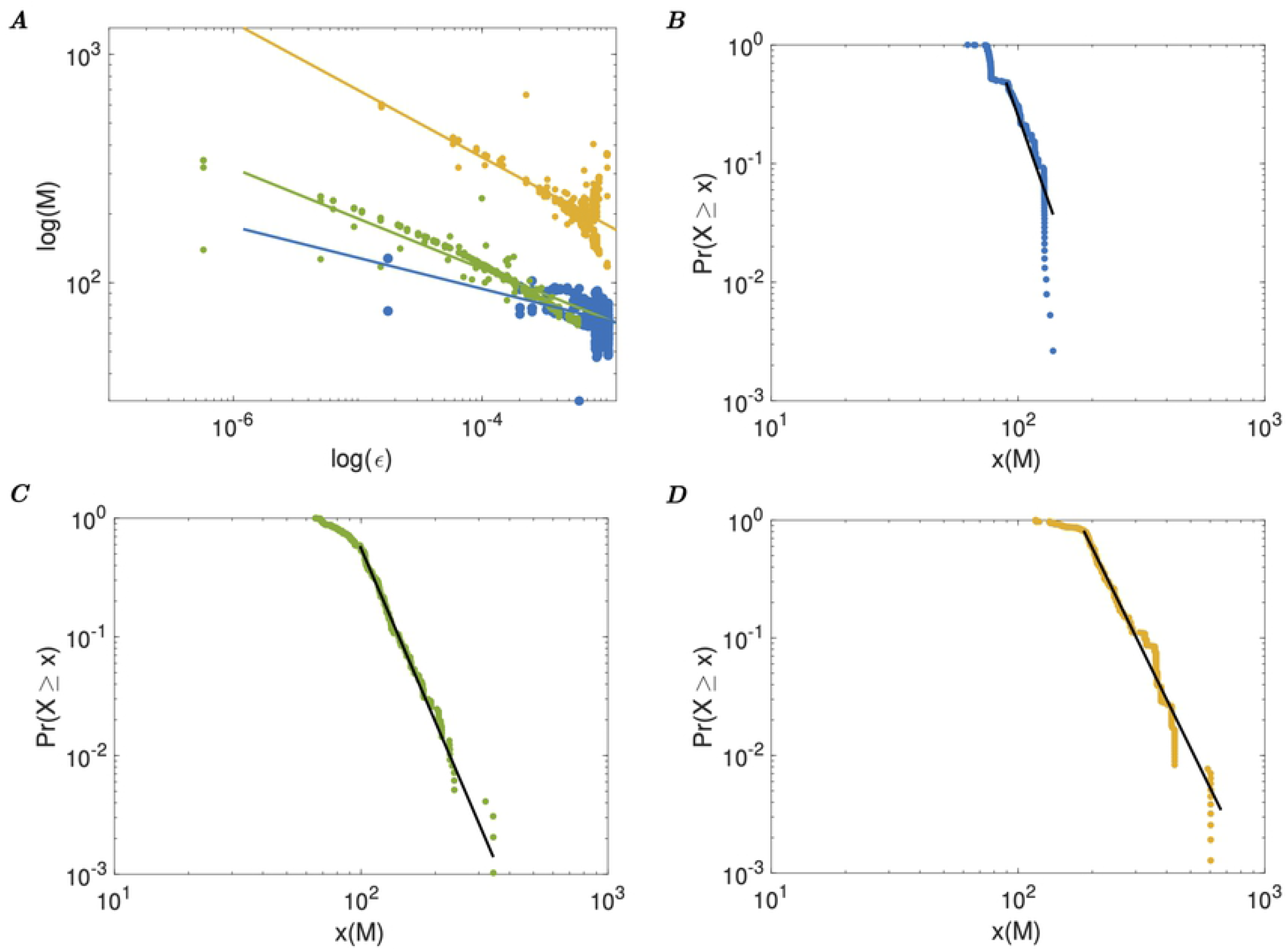
Network size in relation to power law due to random perturbations. ***A*** Log-log plot of the total unweighted sum *M* from 8 (blue), 16 (green), and 32 (gold), weakly coupled RCC controlled systems, with random external perturbations. The power curve fits are *M*(8) = 26.88 *ϵ*^−0.14^, *M*(16) = 15.19 *ϵ*^−0.22^, and *M*(32) = 32.48 *ϵ*^−0.29^. **B** Power-law estimation function for the 8 coupled oscillators with a minimum of *M* = 64.5, and slope *α* = 7.05. **C** Power-law estimation function for the 16 coupled oscillators with a minimum of *M* = 99, and slope *α* = 5.85. ***D*** Power-law estimation function for the 32 coupled oscillators with a minimum of *M* = 184.5, and slope *α* = 5.27.

Using simple finite-difference connections between the oscillators, or more elaborate connectivity schemes as required, the critical dynamics can be generated. The individual oscillators will adjust their local dynamics to accommodate the perturbations by their immediate neighbours. The global dynamics may then become critical due to the nonlinear oscillators. As shown in the result, imposing a linearisation method on the connectivity scheme, such as Crank-Nicholson discretisation, will remove the high frequency nonlinear behaviour and results in the loss of criticality of the global state. The network of oscillating nonlinear models is simulated using standard ODE fixed step integration methods. The fourth-order Runge-Kutta (RK) method, as well as the higher order extensions, Fehlberg RK and Prince-Dormand RK, provide suitable results with varying choice of integration step size. The qualitative results are readily reproducible using any of these methods. The simulation software EuNeurone, used for these models, is available as open source on Github (https://github.com/biodynamical/euneurone). The model equations are available on DataDryad (https://doi.org/10.5061/dryad.hhmgqnkdc) at https://datadryad.org/stash/share/iWjuPsoa_PXLyK0cpOx8JPZUSV1MlNqsGvdiyK0iIaQ

The total unweighted dynamics is determined by the sum of the individual oscillators *i* for a total of *n*, as could be seen by a remote observer for whom the individual oscillators are not readily visible.

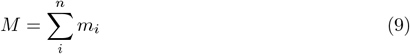

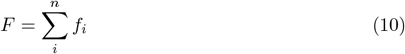

The summed total of the variables *m* for each of the oscillators is added to Eq (8) for the models with global feedback, scaled either negatively or positively as appropriate (*v_k_* = ±0.00001), to ensure that the individual oscillator is within the controlled domain. This feedback is added during the evolution of the differential equations, so that *M* is due to the instantaneous value determined at each integration step, and therefore not merely equal to the current summed input of the previous time step reflecting the local instantaneous state of the system.

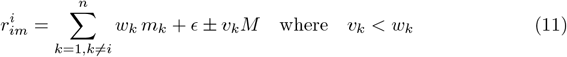

The Eq (11) shows that feedback for this model in effect reduces the connectivity as provided to *r_im_* for each oscillator, but because of the weaker feedback, it does not disrupt the network such as much stronger connectivity or little connectivity would effect.

The log-log plots are generated by the maximal peak values of the unweighted sum max (*M*) during a single oscillation, and used to determine the power plot as estimated using the power-law estimate function according to Clauset et al. [35]. This method estimates the scaling parameter *α* from the power law probability distribution *p*(*x*) = *Pr*(*X* = *x*) = *Cx^α^* using the method of maximum likelihood. This is determined using the numerical maximisation of the logarithmic likelihood function described by

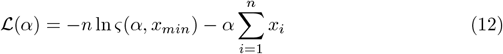

The minimum bound for *x_min_* for these estimates is determined by the Kolmogorov-Smirnov statistic, that is the maximal distance between cumulative distribution functions of the data and the fitted model. It is noted that these methods would require a significantly large data set for reliable estimation. Within the shown simulations, these estimates are primarily to demonstrate that the overall probability of a power law relation is present, but its actual value is not particularly relevant to the argument, given that these models are not at all intended to describe a real physical biosystem as such.

Determining the fitness of any model that can describe the datasets is more readily achievable for probabilistic approaches that allow the preplanned comparisons between models. The goodness of fit, as used above, employing the method of maximum likelihood, is only appropriate to the extent that it allows the model to be reconciled with the options available, in this case the existence of some power law relation. It also suffers somewhat from the ability to be a suitable predictor due to possible overfitting. Similar complexity measures for model estimates are the Bayesian Information Criterion (BIC) and the Akaike Information Criterion (AIC) that can indicate the best fit of log-likelihood, often used for predictors in regression models. These can be determined as follows, based on the Mean Squared Error.

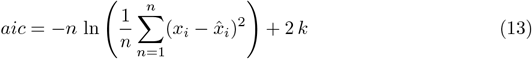

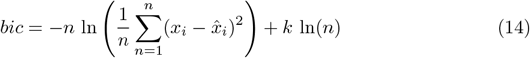

where *k* is the number of parameters of the estimated model, and 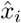 is the estimate fit.

The choice of underlying RCC controlled model is itself also not critical. It has been shown that other RCC controlled oscillators can produce similar critical states, e.g. using the Rössler model [31], but this model has lower adaptability due to the absence of multiple nonlinear terms, preventing the system to respond quickly to the perturbations and nonlinear interactions.

## Results

Any perturbation of the controlled system appears to the RCC method as chaotic change, and it therefore adjusts the rate to compensate. If this is due to noise or an external input, the controlled orbit or state will vary in accordance to the control but remain stable. This allows the construction of networks of controlled nonlinear oscillators that can adjust their dynamic behaviour due to external input. The ability to control dynamics based on local information alone also allows the construction of a mechanism that is capable of control at different scales, due to the fact that each contributing element provides only enough local control to maintain its own stable environment. The cumulative effect, given that these are controlled chaotic elements, would then provide amplification of the control into the global domain. This control of local spatiotemporal dynamics is a further demonstration of interacting rate controlled chaotic systems that exhibit emerging properties which are associated with self-organising criticality with scale-free relations.

### Local control provides global stability

To illustrate this emerging global controlled behaviour, a simulation of 16 coupled RCC controlled chaotic oscillators is used. These oscillators receive local random external perturbations (i.e. external to these oscillators), causing them to change their local dynamics in response. To visualise this, in Fig 1D is shown a short series of only 7 perturbations, where the perturbations are randomly varied every 20,000 time steps. Note that after only very short transients, the total global behaviour of the unweighted sum of the oscillators becomes stable in a different oscillation each time. The phase space plot of a sample of 24 randomly perturbed oscillations is shown in Fig 1E, where the total unweighted sum of the two main variables of each oscillator are plotted, with the transients removed. Each different coloured line represents one of these stable oscillations, showing changes in the dynamics, but only depending on these perturbations. The individual oscillators that combine to generate the global stable dynamics are shown in phase plots in Fig 1F, where each coloured line represents the stable oscillation under the perturbed conditions, with transients removed.

The number of oscillators in the network naturally affects the amplitude of the summed oscillators, but the critical multi-stability property of the network is preserved. This is shown in Fig 2A, where the log-log plot of three networks are shown. The lower graph shows total behaviour *M* of 8 coupled oscillators, the middle of 16 oscillators (same network as shown in figure 1, but with different random perturbations) and on top, 32 coupled oscillators. The curve fits that are superimposed on the three sample behaviours show the power relations *M*(8) = 26.88 *ξ*^−0.14^, *M*(16) = 15.19 *ξ*^−0.22^, and *M*(32) = 32.48 *ξ*^−0.29^. To show that the underlying relation has some power law relation rather than an exponential or similar relation, the power law estimation functions for each of the three samples of coupled networks were determined [35]. In Fig 2B for the 8 coupled oscillators with a minimum of *M* = 64.5, and slope *α* = 7.05. In Fig 2C for the 16 coupled oscillators with a minimum of *M* = 99, and slope *α* = 5.85, and in Fig 2D with a minimum of *M* = 184.5, and slope *α* = 5.27. Due to the limited number of oscillators used, the power law function does not cover many decades. The results show that the near power law relation holds for several sizes of networks; large scale modelling, which is computationally expensive, will need to show the full range of scale-free behaviour.

By modelling a simple network of controlled chaotic nonlinear oscillators, it can be seen that even with random perturbations, the control allows the total system to remain stable. It adapts to these perturbations by stabilising into different orbits. The perturbations can still destabilise the entire system if their contribution is disproportionally large with respect to the ergodic properties of the individual oscillators. The number of elements affects the dynamics by allowing higher amplitudes, but more importantly the system maintains multi-stable states and has apparent power law relations, even for such small networks.

### Scale-free emergent behaviour

To demonstrate how the network of rate controlled oscillators is capable of generating apparent scale-free behaviour, the same network is used, still randomly perturbed by uniform perturbations on each of the oscillators, but simultaneously the network connectivity is stepwise increased. Starting with a relatively low connectivity, strong enough to maintain some cohesion between the oscillators, the connectivity between the oscillators is increased stepwise at a constant rate. This is shown in Fig 3A, where the unweighted sum *M* is shown in time, with stepwise increments of connectivity at each multiple of 10^5^ time steps. Even though the dynamics of the total system may change, it remains stable throughout. Not shown is that if the connectivity becomes too dominant, *M* will explode to a singularity. In Fig 3B is shown the total dynamics of *f* versus *m* of each individual stable oscillation indicated by individual different colours, with transients removed. Although at higher connectivity strengths the orbits are more noisy, due to the amplification of the individual random perturbations, the total system remains stable. This can be seen in Fig 3C, where the sixteen oscillators are shown, for each of the incremental steps, clearly demonstrating their individual stability, even at high connectivity.

**Fig 3.**
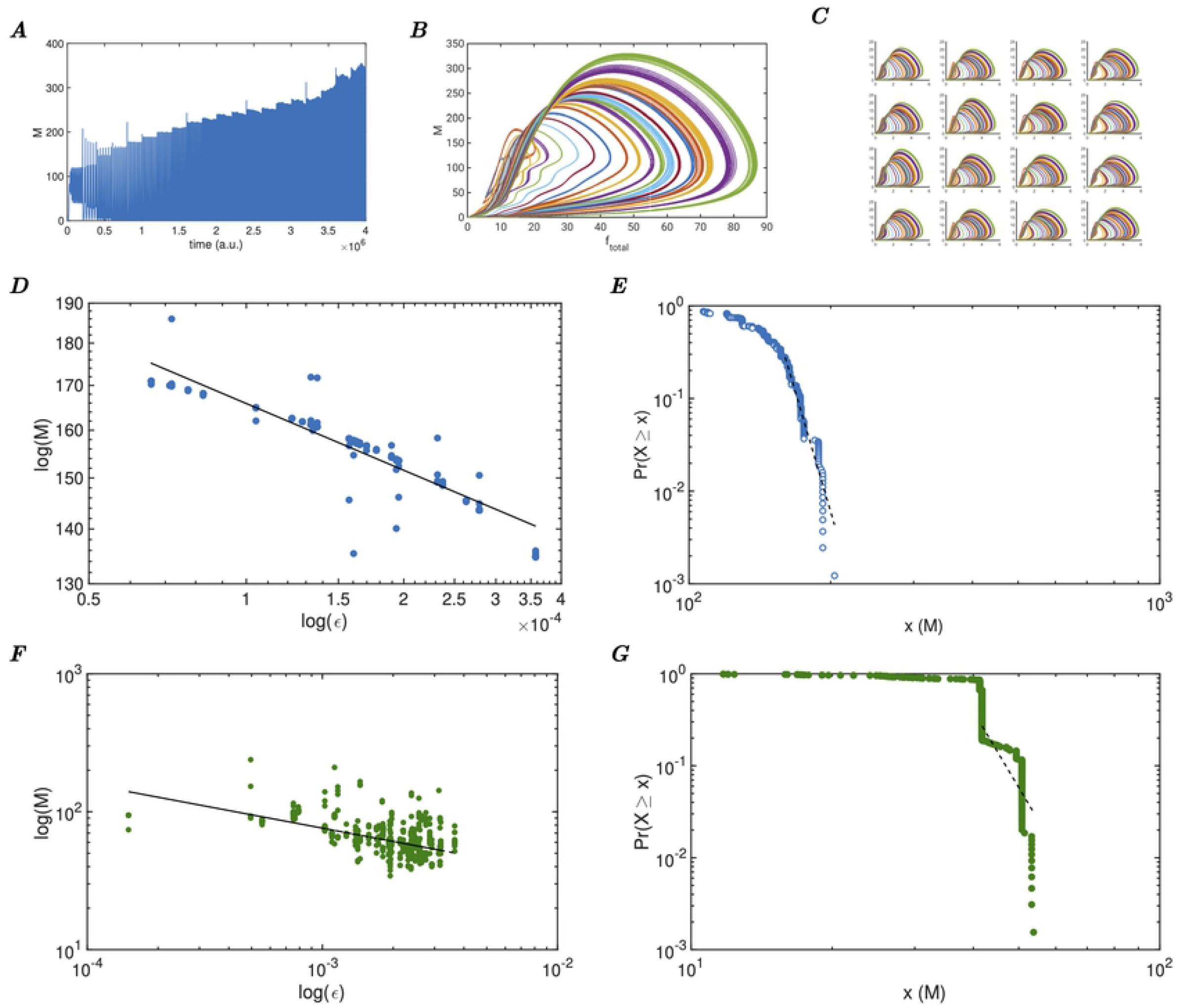
Importance of connectivity in creating an RCC critical system. ***A*** Total unweighted sum *M* from sixteen weakly coupled RCC controlled systems, with random external perturbations, as well as, incremental increases in the connectivity strengths. ***B*** Phase space plot of the total dynamic behaviour of the perturbed systems, with incremental stepwise increases of the connectivity strengths. Each coloured line is one stable oscillation. ***C*** Phase space plots of the sixteen weakly coupled individual systems, with incremental increases in connectivity strengths. ***D*** Log-log plot of the external perturbation e versus the maxima of the total unweighted sum of m. Fitted power curve in black (3.9 *ϵ*^−0.13^). ***E*** Power-law estimation of the maxima of the network with the fitted power function (dotted line) superimposed with minimum *x*(*M*) = 160 with slope *α* = −18.24. ***F*** Log-log plot of the external perturbation *ϵ* versus the maxima of the total unweighted sum *m* with Crank-Nicholson interpolation connectivity scheme. The fitted power curve in black 2.1 *ϵ*^−0.32^. ***G*** Power-law estimation of the maxima of the Crank-Nicholson connected network with the fitted power function (dotted line) superimposed; minimum *x*(*M*) = 41.76, slope *α* = −9.55.

The changes in connectivity could for most coupled dynamic systems be a source of instability, often causing bifurcation phenomena. However, the local control manages to stabilise each individual oscillator, based on local information alone, and thereby ensures that the global system remains stable as well. The subsequent total or global system also demonstrates properties of criticality, in that the total system amplitude grows much faster than merely the sum of the individual elements a linear (or reduced) system would show. To quantify this, the peaks *M* are determined (the maximal value of *M* once stabilised past the transient), and plotted versus the amount of local perturbation in Fig 3D. To demonstrate independence of the connectivity strengths of this scale-free property, the data from the perturbation simulations are used, as in Fig 1D. The log-log plot shows the apparent power-law relation, with a fitted power curve 3.9 *ϵ*^−0.13^. The corresponding power-law estimation function is in Fig 3E, with a minimum of *M* = 160, and slope *α* = 18.24, demonstrating that the data is partly power distributed. It is interesting to observe that this scale-free aspect results in an almost power-law relation which is characteristic for biological observations [15]. Testing the hypothesis that the emergent critical system is based on the combination of the RCC controlled chaotic system with weak to moderately strong connectivity, the connectivity function was modified to match the standard Crank-Nicholson interpolation method, using a two-by-two stencil of the local neighbourhood. The peaks of *M* were determined based on the uniform perturbed network of oscillators, and plotted against the perturbation in Fig 3F. No system parameters were modified, apart from the interpolation connectivity; therefore both the individual oscillators and the total system remained stable, but have lost the scale-free power relation. The power-law estimation function for those data samples is in Fig 3G, with a minimum of *M* = 41.76, and slope *α* = 9.55, showing no power law relation present.

To further investigate the effect of perturbations on a larger pool of oscillators, a network of 64 oscillators was individually perturbed with Normalised Gaussian Distributed perturbations with mean of 0.00005 and variance of 10, to ensure that each oscillator is receiving sufficiently different input. The choice of Gaussian distribution to replace the uniform distribution is made to show that the approach is independent of specific distributions for criticality to emerge. The resulting stable oscillations of 100 orbits are shown in a phase space representation in Fig 4A, where each colour represents a different stable oscillation of the total behaviour of the summed variables *F* versus *M*. In Fig 4B can be seen the maxima of each stable oscillation of *F* versus *M*, showing that there are two distinct domains of oscillatory behaviour emerging from this network. The first type is a single orbit that seems almost linearly related. The second type is a two-orbit that causes an apparent cluster on the left, above the apparent line. However, the underlying relation between the perturbations and the effect on the total dynamic behaviour is not linear, as can be seen in Fig 4C, where the total summed perturbations of the 64 oscillators is plotted against the maxima of *M*. It becomes clear that the left part of this plot for *ϵ* <= 0.043 (before the dotted line), exhibits a power-law relation between the total perturbation *ϵ* and the maxima of *M*. This is shown in a log-log plot in Fig 4D, where superimposed are also shown four curve fits. The power function 134.7 *ϵ*^−0.328^ in red is fitted to the curve for *ϵ* <= 0.043 and clearly expresses the sub-domain where a power law relation exits. In blue is shown the fit of an exponential function 715.9*e*^−16.89ϵ^ on the same domain, which is clearly quite poor. The domain for *ϵ* > 0.043 fits a power law 5.38*e* + 05 *ϵ*^2.254^ in orange) but also an exponential function in green 56.59*e*^48.26ϵ^, indicating that this domain does not have a power law relation, and may be exponential or simply linear.

**Fig 4.**
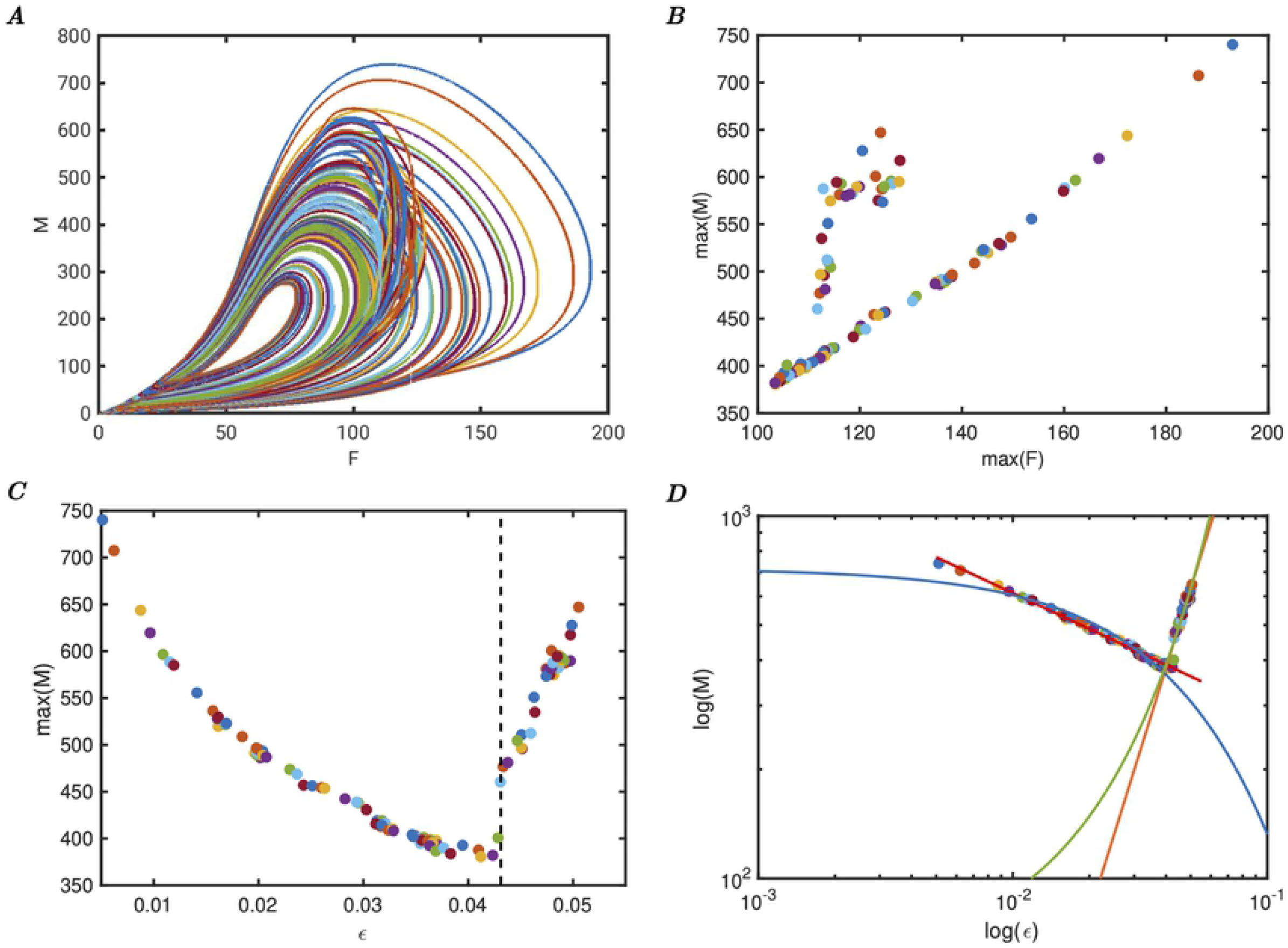
Domains of power law relations within perturbation space of RCC controlled Self-Organised Criticality. ***A*** Phase space plot of *M* versus *F* of 100 stable oscillations in a 64 oscillator network. ***B*** Plot of the maxima of *F* versus the maxima of *M* of the 100 orbits on the left. Notice the two domains where the dynamics of the oscillators change. ***C*** Plot of the Normalised Gaussian distributed perturbation *ϵ* with variance 10 versus the maxima of *M*, showing that around 0.043 the oscillators change their shape. ***D*** Log-log plot of the Normalised Gaussian distributed perturbation *ϵ* versus the maxima *M*, same as the panel on the left. Additionally, the curve fits for the power functions 134.7 *ϵ*^−0.328^ (red), 5.38 × 10^5^ *ϵ*^2.254^ (orange), and exponential function fits 715.9*e*^−16.89ϵ^ (blue), 56.59*e*^48-26ϵ^ (green). This shows clearly that the system can have a power law relation for limited size perturbations, and a non-power law relation otherwise.

These models are based on deterministic behaviour of the perturbed network which itself is based on controlled chaotic oscillators. The models are only perturbed from one state to another by the random perturbations, and no noise is included in these systems. The resulting behaviour is therefore fully stable deterministic and is completely described by the RCC control. It is possible to determine the log-likelihood of the data for model selection. The relevance of probabilistic model selection approaches is very limited to this type of modelling because statistics such as AIC are designed for preplanned model estimates that do not take into account the model parameter space, which for a large set of nonlinear differential equations is extensive [36, page 216]. It should also be considered that the underlying model is not based on probabilistic methods, and that the emerging power or exponential relation causes great variance in the data, which greatly amplifies the mean squared error. As a representative example of such estimates, the BIC and AIC were determined for both the power fit model, and the exponential fit model on the data shown in Fig 4**D**.

The model fits with samples for *ϵ* < 0.043 are 135.5 *ϵ*^−0.3277^, and 712.7*e*^−16.65ϵ^. In this case, for the power fit, the Mean Square Error is 46.2, BIC is −247.98., AIC is −254.64; and for the exponential fit MSE is 374.74, BIC is −390.33, AIC is −396.98. Despite the relatively large value of MSE, the power fit still seems better suited than the exponential fit. For the model fits with samples *ϵ* > 0.043, the fits are 5.387*e* + 5 *ϵ*^2.254^, and 56.59*e*^48.26ϵ^. The corresponding values for the power fit, MSE is 304.47, BIC is −183.85, AIC is −188.43; and for the exponential fit, MSE is 330.63, BIC is −186.65, AIC is −191.23, which shows there is little difference in these estimates for model selection, possibly favouring the power fit somewhat.

It can also be concluded from this set of simulations that the relation between perturbations, the number of oscillators, and connectivity strengths is not straightforward. However, the resulting system of connected oscillators is dynamically stable, can exhibit different types of emerging relations between the total amplitude of the oscillators and the perturbations, and also that the emerging power law relations are an epiphenomenon of the network’s attempt to stabilise its overall behaviour in response to external random perturbations.

These apparent power-law relations provide a strong indication that the nonlinear behaviour of each oscillator causes perturbations such that the total behaviour of these oscillators is much stronger than the individual contributions they provide. This does not seem to be based on the onset of synchronisation due to changing coupling strengths, because it occurs for constant coupling and there is no critical coupling strength for which the relation holds.

It has already been shown that chaotic units may generate a critical system [37], where the effect of connectivity over time is shown to have a power-law relation between connectivity and the connected chaotic systems. Here, the connectivity level of *k*-connectedness is critical to establish the power law stability. Therefore, a locally controlled mechanism of generating critical rate controlled systems, as described in this paper, may provide the necessary dynamics for complex interactions found in biological systems. For example, local producers of a protein can regulate their behaviour on local information alone, but still provide the effect needed by remote consumers of the protein. This may be particularly important for regulatory processes and may provide the key elements of a feedback loop process that responds rapidly to changing global behaviours.

### Criticality as a homeostatic process

Adapting the network of sixteen coupled oscillators to include a global feedback of the total dynamic behaviour is possible by adding the scaled total behaviour to the external perturbation of each oscillator, ensuring that the perturbed input to each of the oscillators is not too great to push it out of the controlled domain. By making this feedback either positive or negative, the effect of global input to the individual oscillators as negative and positive feedback loops is shown. In Fig 5A is shown the total dynamics of the network, with stepwise increased connectivity and positive feedback, but without further random perturbations. In this case, the different stable orbits due to the increased connectivity are amplified to higher totals than without the positive feedback. The corresponding power-law estimation function, based on the maximal values of these orbits, shows a strong power relation between the connectivity and the total amplitude of the oscillations (Fig 5B). Conversely, if the feedback of the total behaviour is negative, the total dynamics is more compressed, as can be expected (Fig 5C). Also, the individual oscillations are more limited, with less drift. The power-law estimation function of the negative feedback (Fig 5D) shows, interestingly, that the critical system has lost its power-law relation and appears more linear. This would suggest that, just like in classical homeostatic negative feedback control, the global negative feedback can stabilise individual global states. This in effect counteracts the local connectivity, but at the expense of less dynamic capabilities. Due to the currently available network connectivity the network has become less complex.

**Fig 5.**
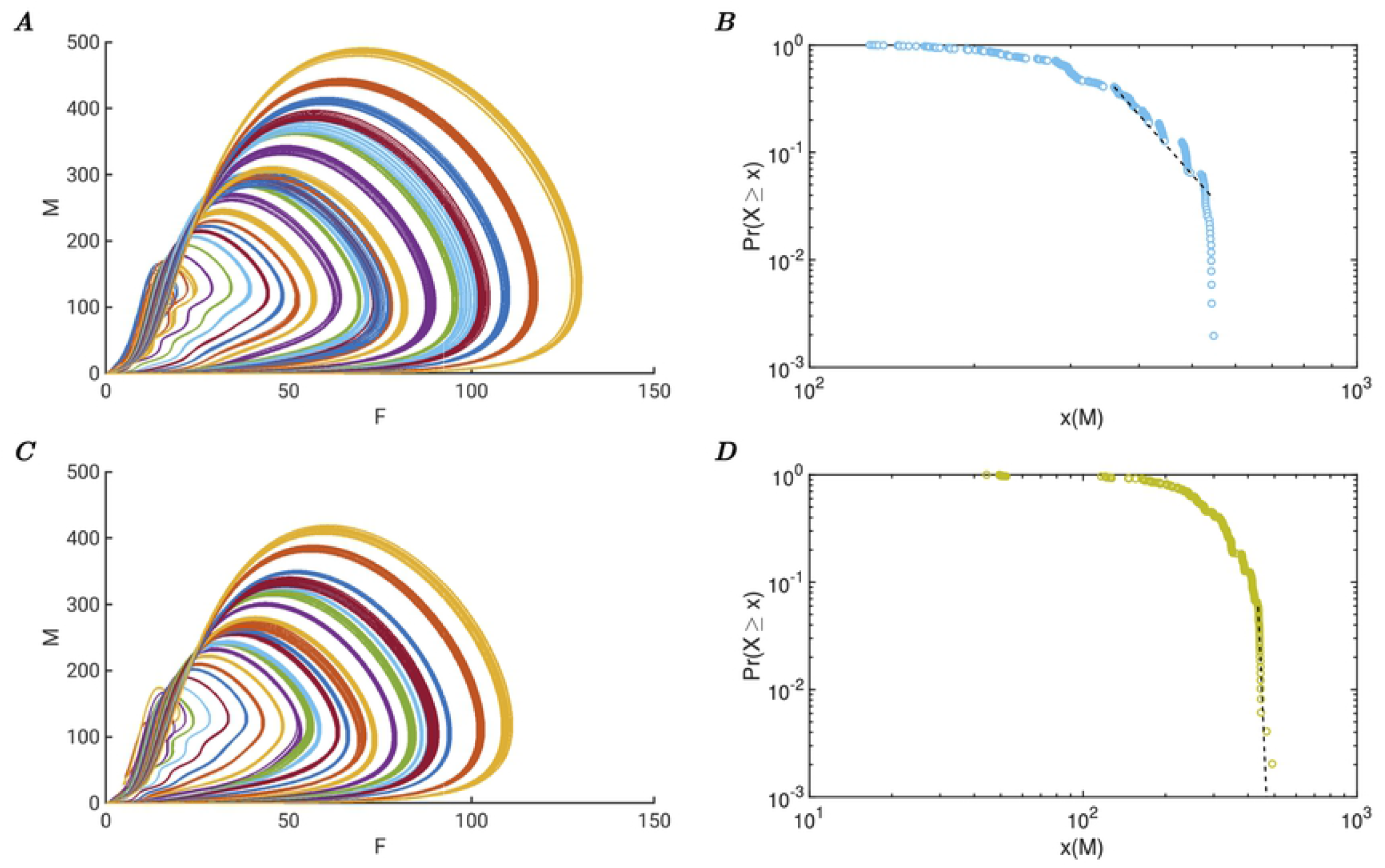
Global positive or negative feedback causing enhancement or loss of power relations. ***A*** Total dynamic behaviour of a network of sixteen coupled rate controlled oscillators with global positive feedback. The feedback pushes the total dynamic behaviour to new heights. Each coloured line is one stable oscillation with positive feedback. ***B*** Power-law estimation of the maxima of the positive feedback network with the fitted power function (dotted line) superimposed with minimum *x*(*M*) = 360.24 and slope *α* = –6.74. ***C*** Total dynamic behaviour of a network of sixteen coupled rate controlled oscillators, differently coloured lines indicate different stable oscillations, with negative global feedback. The negative feedback reduces the behaviour. ***D*** Power-law estimation of the maxima of the negative feedback network with the fitted power function (dotted line) superimposed with minimum *x*(*M*) = 435.69 and slope *α* = –59.29.

## Conclusion

Self-Organised Criticality emerges from local nonlinear interactions of Rate Control of Chaos controlled elements. This allows critical states to exist where the global dynamics, as expressed by the total unweighted sum of each element, is dynamically stable. Multiple states are achievable for the dynamic system, driving the system towards the desired global state by perturbing individual oscillators. Adjusting the local connectivity, in effect recruiting more elements, allows different dynamic states to emerge. Increasing the number of elements allows domains of different relations between perturbations and total behaviour to emerge. These apparent relations, whether power-law, exponential or linear relations, appear due to the networks’ self-stabilising properties. Given that he primary aim of any biological system is to maintain stability, the exact nature of the emerging relation becomes therefore a mere epiphenomenon of the ability of the system to stabilise and control its complex dynamics. Furthermore, providing localised global feedback may allow other critical states to also become available, or allow the control of global behaviour into a more limited stable domain.

The described models are clearly in a critical dynamic state, and can readily change state due to local perturbations. This critical state is the result from the RCC controlled systems with local interaction between the units and is therefore self-organising. Lastly, the emerging power law relations and other relations are the result of the nonlinear interactions of the oscillatory units. Therefore, these models are said to describe Self-Organising Criticality because it matches the characteristic key features: non-trivial scaling, spatiotemporal power-law relations (domain bound), and self tuning to the critical state [7].

Biosystems can therefore emerge from the localised interactions between controlled nonlinear systems, creating the perfect combination of complexities that supersedes the limitations of linear systems, avoids the instability of chaotic nonlinear systems, and limits the domain of self-emergent critical systems. This opens up the possibility of innovative research in controlled nonlinear biological dynamics with direct applications to health, engineering control, and human wellbeing.

